# Auto-regressive modeling and diagnostics for qPCR amplification

**DOI:** 10.1101/665596

**Authors:** Benjamin Hsu, Valeriia Sherina, Matthew N. McCall

## Abstract

Current methods used to analyze real-time quantitative polymerase chain reaction (qPCR) data exhibit systematic deviations from the assumed model over the progression of the reaction. Slight variations in the amount of the initial target molecule or in early amplifications are likely responsible for these deviations. Commonly-used 4- and 5-parameter sigmoidal models appear to be particularly susceptible to this issue, often displaying patterns of autocorrelation in the residuals. The presence of this phenomenon, even for technical replicates, suggests that these parametric models may be misspecified. Specifically, they do not account for the sequential dependent nature of qPCR fluorescence measurements. We demonstrate that a Smooth Transition Autoregressive (STAR) model addresses this limitation by explicitly modeling the dependence between cycles and the gradual transition between amplification regimes. In summary, application of a STAR model to qPCR amplification data improves model fit and reduces autocorrelation in the residuals.

## Introduction

### 1.1 Background

Polymerase chain reaction (PCR) is a molecular biology technique used to amplify the copies of a specific DNA sequence. Real-time polymerase chain reaction, also known as quantitative polymerase chain reaction (qPCR), is a widely applied laboratory technique based on PCR to allow for gene quantification. Despite the advent of alternative technologies to measure gene expression, e.g. microarrays and high-throughput sequencing, qPCR remains one of the most extensively used methods for targeted RNA quantification. This is in part due to its accuracy and sensitivity to small transcriptional changes, in addition to comparably lower cost.

Quantitative PCR amplification curves typically have multiple stages: a baseline phase in which fluorescence remains approximately constant, an exponential phase in which the fluorescence approximately doubles for each cycle, and a plateau phase as the amplification tapers off (Figure 1). During the baseline phase, there is a slow increase in amplicon product but the corresponding increase in fluorescence is masked by a substantial amount of background noise. One way to reduce background noise is using baseline subtraction methods; the simplest of which is subtracting the minimum observed fluorescence from each fluorescence measurement. This eliminates some variability between reactions but does not address the fundamental lack of a detectable fluorescence signal. The next phase occurs when the signal gets strong enough to separate from the background noise. The reaction enters the exponential phase where the amplification reaches peak production. Then, as reagents are consumed and the reaction is limited by the number of available nucleotides, the amplification slows down and plateaus, concluding the PCR reaction.

**Figure 1.**
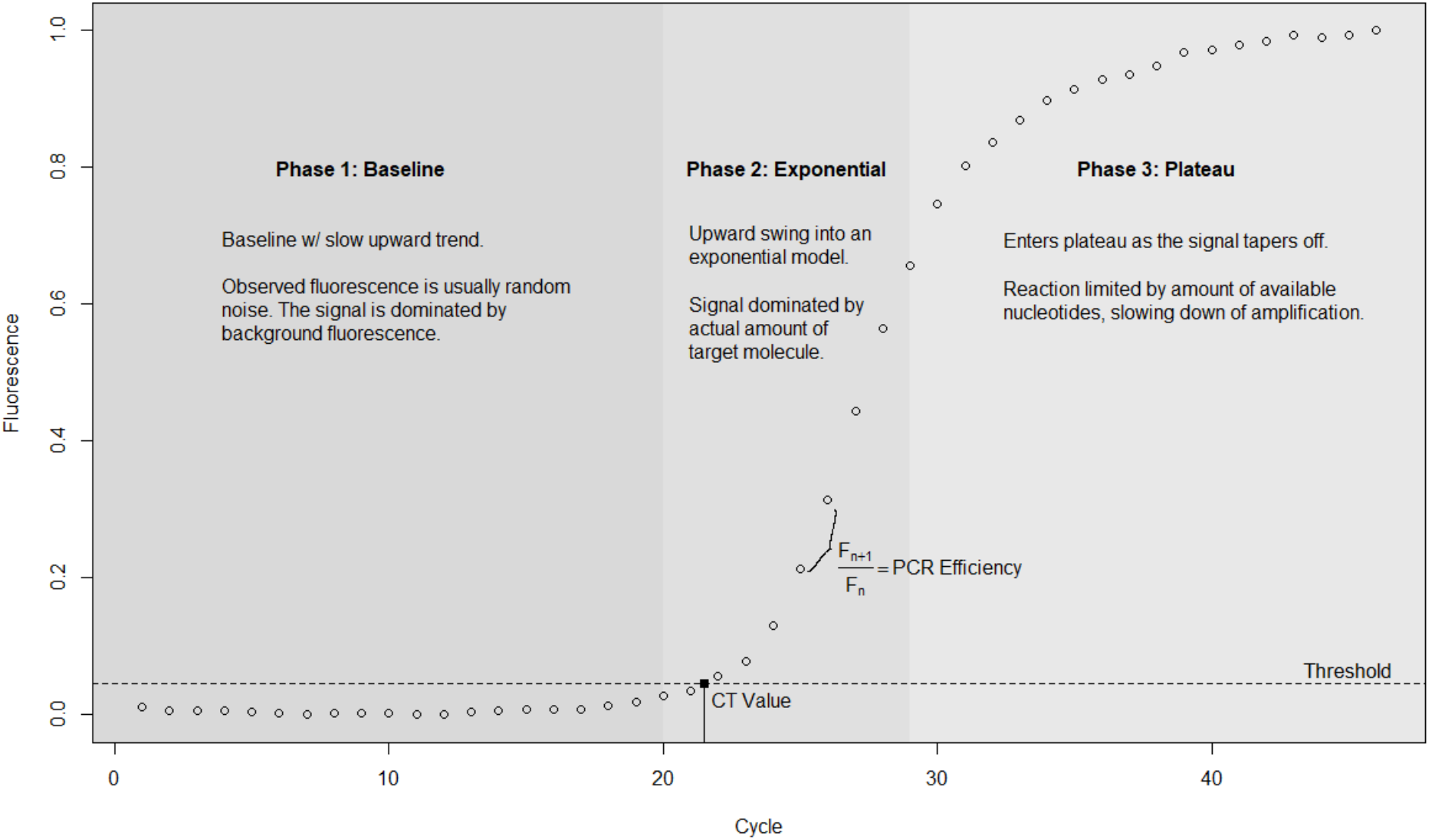
The amplification curve of hsa-miR-520e_001119 (replicate E1) in the miRcomp data set. The dashed horizontal line is the threshold above the background fluorescence. The CT value is the black square. PCR efficiency is calculated by taking the fluorescence value in the later cycle divided by the former cycle’s value. There are three phases shown: (1) baseline, (2) exponential, and (3) plateau.

Historically, the entire amplification curve is summarized by the cycle at which the fluorescence exceeds a threshold, known as the cycle threshold (CT) value. A smaller CT value means that it took fewer cycles for the fluorescence to cross the threshold, meaning the target molecule was more abundant in the initial sample. Earlier methods relied on manually selecting a threshold slightly above the amplification baseline; however, this required visual inspection of each amplification curve which is infeasible for larger experiments. This led to the development of automated threshold selection methods as well as alternative procedures to quantify expression from qPCR amplification data. These latter methods often produce cycle quantification (Cq) values on the same scale as the CT values but without the use of a threshold (Bustin et al. 2009).

Many of the most widely used methods to estimate expression from qPCR data rely on fitting a 4- or 5-parameter sigmoidal curve (Spiess et al. 2008). The fitted sigmoidal curves are then used to estimate Cq values via the second derivative method (SDM), which finds the cycle at which the second derivative is maximized. This corresponds to the point in the PCR reaction with the sharpest increase in fluorescence which typically occurs at the start of the exponential phase (Spiess et al. 2008). SDM allow for high-throughput quantification of qPCR amplification data.

In addition to estimating Cq values, the amplification data can be used to estimate amplification efficiency, which provides information about the rate at which the concentration of the target molecule is increasing as the reaction progresses. A simple generative model describes the relationship between observed fluorescence, the initial number of target molecules, and the PCR efficiency. The number of target molecules at cycle T is denoted as N_T_. This is equivalent to the initial number of molecules (at cycle 0), n_0_, multiplied by the product sequence of the efficiency of the PCR reaction at cycle t, r_t_. The fluorescence at cycle T (F_T_) is assumed to be a monotonically increasing function of the number of molecules plus measurement error.

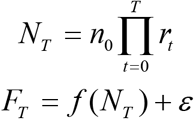

Theoretically, the efficiency should be equal to 2, representing a doubling of the target molecule at each PCR cycle (Livak and Schmittgen 2001). However, PCR efficiency differs from cycle to cycle, and any Cq estimate under the assumption of static efficiency is likely to be unreliable (Bar et. al 2011). In addition, inhibited reactions that delay when the cycle reaches the threshold also result in underestimation of the initial target abundance (Lievens et. al 2011). Because small errors in modeling qPCR amplification data can produce significant changes in subsequent estimates, it is crucial to minimize these errors in the initial modeling.

### 1.2 Sigmoidal Models

4 and 5-parameter sigmoidal models use a non-linear fit to provide better estimation of threshold fluorescence, cycle-to-cycle efficiency approximations, and target quantification. Both 4- and 5-parameter sigmoidal models contain parameters corresponding to lower and upper asymptotes, slope, and inflection point (Spiess et al. 2008). The 5-parameter sigmoidal model introduces an additional asymmetry parameter that allows for differences in the curvature before and after the inflection point. By considering asymmetry, the estimation of the fluorescence and cycle threshold in the exponential phase is improved, thereby, yielding more accurate quantification of the target molecule.

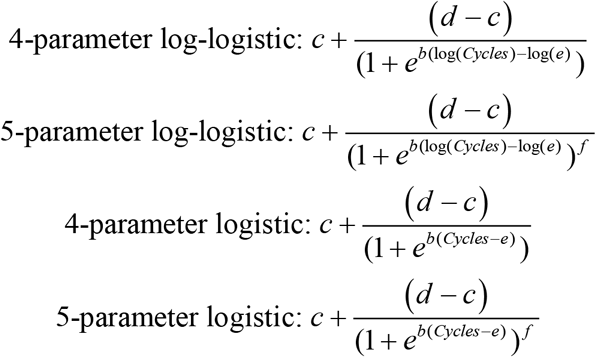

Although the 4-parameter and 5-parameter sigmoidal models approximate the typically structure of a qPCR amplification curve, these models do not capture the sequential dependence of the amplification process. Mechanistically, the amount of the target molecule, and thereby the observed fluorescence, at a given cycle depends on the amount at the previous cycle, but under the 4- and 5-parameter models, residual variation at each cycle is considered independent. Despite this limitation, the sigmoidal models still provide the ability to capture the general shape of the curve, and more importantly, efficiently analyze amplification curves while increasing reproducibility in qPCR analyses (Rutledge 2011). For these reasons, we will compare the performance of our proposed model to these sigmoidal models.

### 1.3 Smooth Transition Autoregressive (STAR) Model

We propose using autoregressive time series models to model qPCR amplification data because the fluorescence value at a given cycles depends on the previous cycles. Note that a simple autoregressive model, in which the fluorescence in the n^th^ cycle is linearly dependent on its own previous values cannot effectively model the amplification process, since the relationship between qPCR cycle and fluorescence changes between specific phases of the reaction (Supplementary Figure 1). Different phases in the amplification process are characterized with their own state-dependent behavior. A single autoregressive model would fail to capture this behavior.

Threshold Autoregressive (TAR) models are constituted by discontinuities that allow for a switch in regime once the threshold is exceeded. When the regimes are governed by the lagged time series itself, the model is a Self-Exciting TAR (SETAR) model. The SETAR model is set up with autoregressive processes, AR(p), and an indicator function for event A, I[A], based on the threshold variable, q_t_, and the bordered threshold, c (Tong 1978, Lim and Tong 1980, Tong 1990). Here, we show a TAR model with 2 regimes, with a lag of 1, threshold c, and threshold variable of the 1 time lag of itself. We specify an AR(1) model for both regimes, and produce the 2 regime SETAR model:

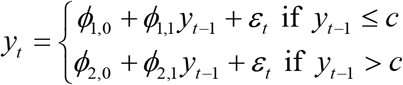

This can be re-written with indicator variables as:

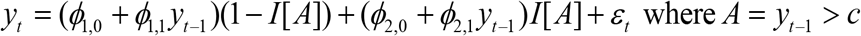

Unlike discontinuities in TAR models where each subsection follows different AR(p) models, PCR data is a continuous process that is better modeled by a gradual transition between regimes. Thus, we chose to apply a Smooth Transition Autoregressive (STAR) model that models the regime changes by a continuous function. The STAR model is similar to the SETAR model, with the exception that the indicator function, I[A], is replaced with the transition function G(-) that monotonically increases from 0 to 1 (Terasvirta 1994). The addition of the smoothing parameter, gamma, will specify how abruptly the switch between the two regimes occurs at q_t_=c. For the purpose of this paper, we will focus on the logistic function as our transition function and refer to the model as a Logistic Smooth Transition Autoregressive (LSTAR) model. In this paper, we consider the following class of two regime LSTAR models:

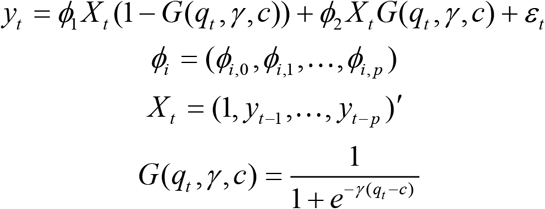

In contrast to the sigmoidal models explored above, the STAR model allows for a flexible transition between different stages in the baseline and upper phases of amplification curves through the use of a logistic smoothing transition function.

### 1.5 Aim of the Study

We evaluate the performance of both sigmoidal and autoregressive models on two benchmark data sets. Our analysis starts with a visual examination of the residuals from each fitted model. Any underlying residual trends may result in biased gene quantification. This is followed by an examination of several measures of autocorrelation (e.g. Durbin-Watson, Ljung-Box, and Pearson), all of which assess aspects of the models’ residuals over time. An ideal model should exhibit no residual auto-correlation. Beyond assessing residual patterns, we calculate the residual sum of squares (RSS) as a translatable measurement of performance amongst fitted models. We propose a more informative measurement model fit by focusing on the RSS within 2 cycles of the Cq value, which we call the local RSS. We focus on this region because it has the most impact on estimation of the Cq value.

## Methods

### 2.1 qPCR Datasets

We used two previously published data sets. The first is a 96-well reaction replicate dataset of Glycerinaldehy-3-phosphat-Dehydrogenase (GAPDH) amplified with the Life Technologies StepOne Plus real-time PCR system for 40 cycles, which is described in more detail in Spiess (2008). This data set will be referred to as GAPDH.SO. The GAPDH.SO data are comprised of of 8 subsets each consisting of 12 replicates. The fluorescence values in this data set were not baseline subtracted.

The second data set used is a qPCR-based microRNA array that consists of 754 human microRNAs across two sets of primer pools, which is described in more detail in McCall et. al (2016). This data set will be referred to as miRcomp. The cycle length for the reactions are either 40 or 46 cycles, depending on the target and replicate. The miRcomp data consists of 10 mixture / dilution conditions with 4 replicates each. Unless otherwise specified, the fluorescence values used from this data set will be baseline subtracted.

### 2.2 Implementation

The sigmoidal models were fit using the qpcR R package, and the Cq value was estimated as the cycle at which the second derivative of the fitted curve was maximized, referred to as cpD2 in the qpcR package. The LSTAR model was fit using the tsDyn R package. The expression estimates are defined as the midpoint between the two regimes, i.e. the cycle at which the transition function equals 0.5. All statistical tests were carried out in the R open source programming environment using standard functions.

### 2.3 An LSTAR Model for qPCR

To model qPCR dynamics, we adopt a nonlinear time series model with each regime explained by a linear autoregressive model. This captures key attributes of PCR dynamics. First, PCR reactions tend to consist of distinct regimes. Second, fluorescence increases with the number of cycles. Third, there should be a gradual change from one regime to the next, despite an independent autoregressive model used in each state. To satisfy these criteria, we propose the following Smooth Transition Autoregressive Model with a logistic transition function:

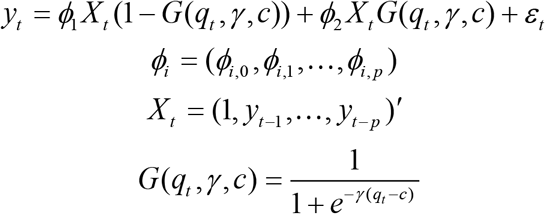

To fit an LSTAR model, we have to decide on the threshold variable, threshold value, smoothness parameter, lag-term, and embedding dimension. The threshold variable, qt, is responsible for gradual change in the transition function, and the threshold value, c, is the point of equality between the lower and upper regimes. Meanwhile, the smoothness parameter, γ, specifies how abrupt the transition is at q_t_=c. The independent autoregressive models of each regime are specified by the number of lags and m-dimensions included. The lag-term and embedding dimension encode the dependence on the previous cycle information and cyclical trends, respectively. Since there are no cyclical trends in qPCR data, we omit any cyclical multiplier. To do so, we restrict the lag-term to 1, and alter the m-dimension to capture all lags that are multiples of 1 (see Supplementary Note). In addition, we require the lower and upper regimes to have the same number of lags. This will produce autoregressive models of the same form for each regime, with different coefficients. For qPCR data, we set the lag-term to either 1 or 2 because the AIC decreased substantially less with the addition of lag terms greater than 2 (Supplementary Figure 2). Additionally, because qPCR amplification processes are typically 40 cycles, inclusions of lags over 2 would truncate the data set with little benefit. Finally, we specify the threshold variable to depend on the current cycle or previous cycle (lag 0 or lag 1). The threshold value and smoothness parameter are estimated by using the Broyden-Fletcher-Goldfarb-Shanno (BFGS) optimization algorithm.

### 2.4 Assessments of Model Performance

#### 2.4.1 Model Abbreviations

The methods are abbreviated as follows:

- 4-parameter log-logistic sigmoidal model (l4)
- 5-parameter log-logistic sigmoidal model (l5)
- 4-parameter logistic sigmoidal model (b4)
- 5-parameter log-logistic sigmoidal model (b5)
- logistic smooth transition autoregressive model (LSTAR)

#### 2.4.2 Residuals and Local Residuals

For both the GAPDH.SO and miRcomp data sets, we evaluate model fit by the residuals sum of squares (RSS) and the residuals sum of squares around the Cq values (local RSS). The RSS is the sum of all the residuals squared, while the Local RSS is the sum of the residuals squared for residuals within 2 cycles of the Cq values. The model fit in that region has the most impact on estimation of the Cq value.

#### 2.4.3 Autocorrelation of Residuals

Residuals are expected to be mean zero, constant variance, and uncorrelated. The presence of autocorrelation in the residuals may be indicative of a model that fails to accurately model PCR dynamics. We use Durbin-Watson, Ljung-Box, and the lag-1 Pearson correlation to test for autocorrelation amongst the residuals.

The null hypothesis of the Durbin-Watson test is that the residuals are not serially correlated, which is tested against the alternative that the residuals can be explained by a first order autoregressive model. Specifically, the Durbin-Watson statistic tests whether the residuals to follow a white noise process, which is a specific stationary process where the residuals have mean zero, constant variance σ^2^, and zero autocovariance except at lag zero. The test statistic can be approximated using residuals from any model (see Chen 2016) as follows:

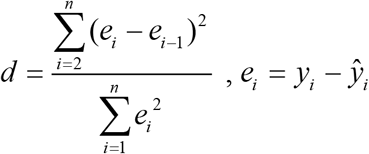

When the autocorrelation coefficient is equal to 0, the Durbin-Watson statistic is equal to 2; therefore, it is assumed that under this scenario the residuals are serially correlated. When the autocorrelation coefficient is close to 1, this suggests perfect positive correlation in the residuals as the Durbin-Watson test statistic approaches 0. Autocorrelation close to −1 would denote perfect negative correlation in residuals and the Durbin-Watson test statistic would approach 4.

The Ljung-Box test (Ljung and Box 1978) considers the null hypothesis that the residuals are random with a correlation of zero. The test statistic is given below, where n is the sample size, k is the lag term, m is the number of lags tested, and rho_k is the autocorrelation at lag = k.

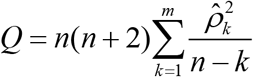

The test statistic follows a chi-squared distribution with m degrees of freedom.

Finally, the lag 1 Pearson correlation, defined as the correlation between the residuals and the one-term lagged residuals, provides an easily interpretable summary of the autocorrelation.

## Results

### 3.1 Residual Patterns

We applied the four sigmoidal models to the GAPDH.SO data set in two ways: (1) to estimate an average curve for all eight technical replicates and (2) to estimate a separate curve for each technical replicate. First, a subset of GAPDH.SO data with replicates was analyzed using a weighted nonlinear least-squares fitting algorithm, also known as the Levenberg-Marquardt algorithm, which estimates a single model based on the replicate values. When this one fitted curve is used to represent a group of replicates and the residuals are computed between the average curve and the observed values, there is a consistent replicate-specific bias (Figure 2 top row). Additionally, the magnitude of this bias appears to increase starting at the beginning of the exponential phase of the reaction, slightly before the Cq value estimate (vertical line in Figure 2). Even after base-line subtraction, we see similar patterns in the model residuals (Figure 2 bottom row). In the first few cycles, the residuals are now close to 0; however, the same biases appear at the start of the exponential phase. Thus, the baseline subtraction method does not appear to alleviate this effect. Similar patterns can be seen in the miRcomp data set (Supplementary Figure 3).

**Figure 2.**
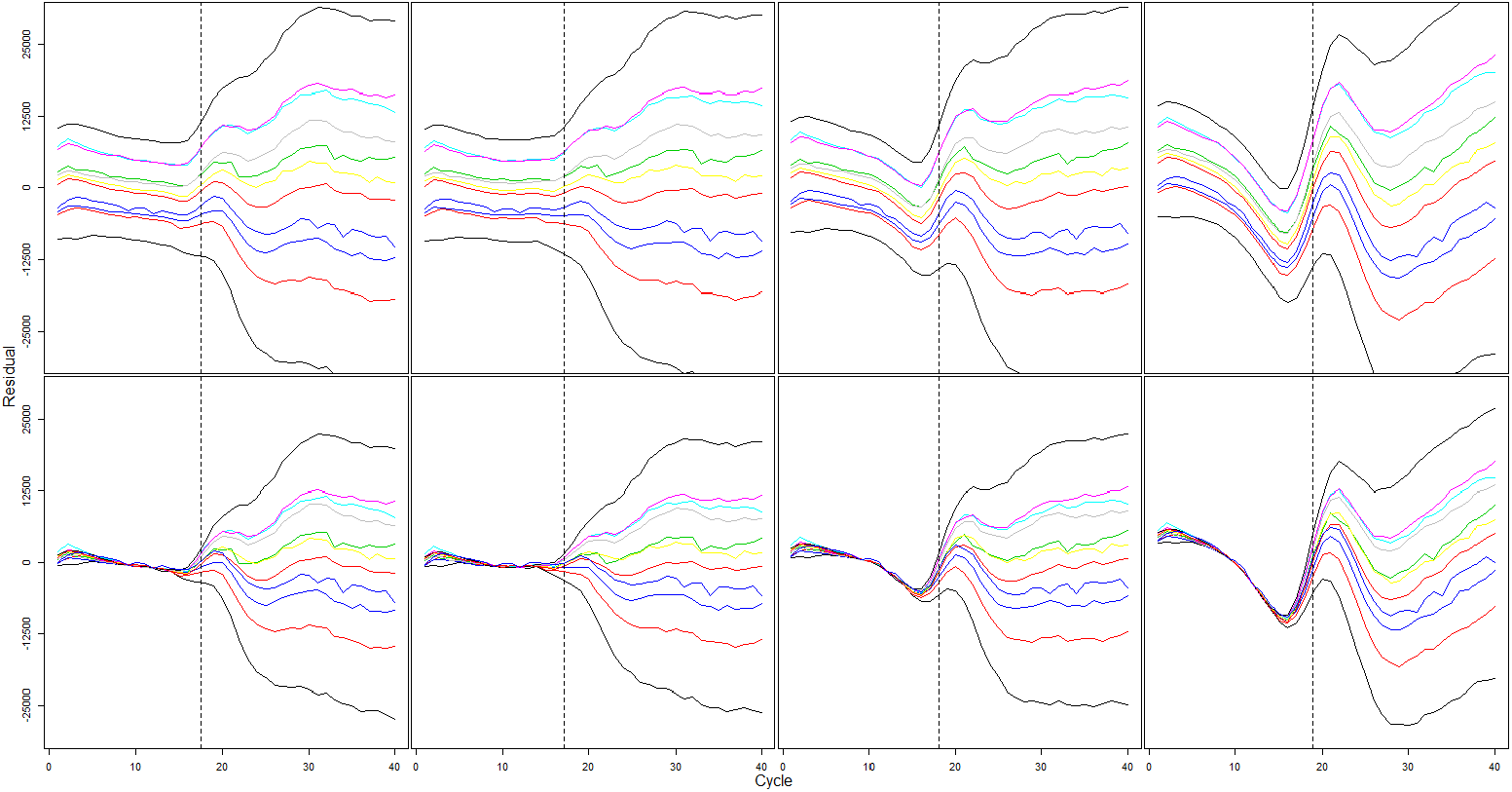
(Left to right: 5-parameter log-logistic model (l5), 5-parameter logistic model (b5), 4-parameter log-logistic model (l4), 4-parameter logistic model (b4). Top row: raw fluorescence, bottom row: baseline subtracted fluorescence) The Levenberg-Marquardt algorithm was used to estimate one representative curve for all 8 technical replicates. The residuals for the representative curve vs. replicates within the subset are plotted by cycle progression using the specified sigmoidal model for one set of replicates in the GAPDH.SO data set. The dashed vertical lines are the Cq values of the representative curve for each subset using the second derivative method (SDM).

Next, instead of estimating one representative curve for each set of replicates, we estimate an amplification curve for each replicate. The residuals from these fitted curves show a different type of pattern than those from the average fitted curve. In Figure 3, the cycle-to-cycle residuals show little variation amongst each of the replications within a subset. The observed patterns of residual bias suggest a systematic lack of model fit. The consistency in this bias between replicates demonstrates the reproducibility of the observed systematic deviation from the sigmoidal models. Similar to the between replicate residuals, transitions into and out of the exponential phase, primarily in cycles 10 to 20, and 20 to 30, respectively, appear to result in the largest bias. In addition, we the ranges of the residuals is smaller for 5-parameter models compared to the 4-parameter models. This supports the use of 5-parameter models as they seem to provide a better fit, but similar patterns of bias persist. As before, similar patterns can be seen in the miRcomp data set (Supplementary Figure 4).

**Figure 3.**
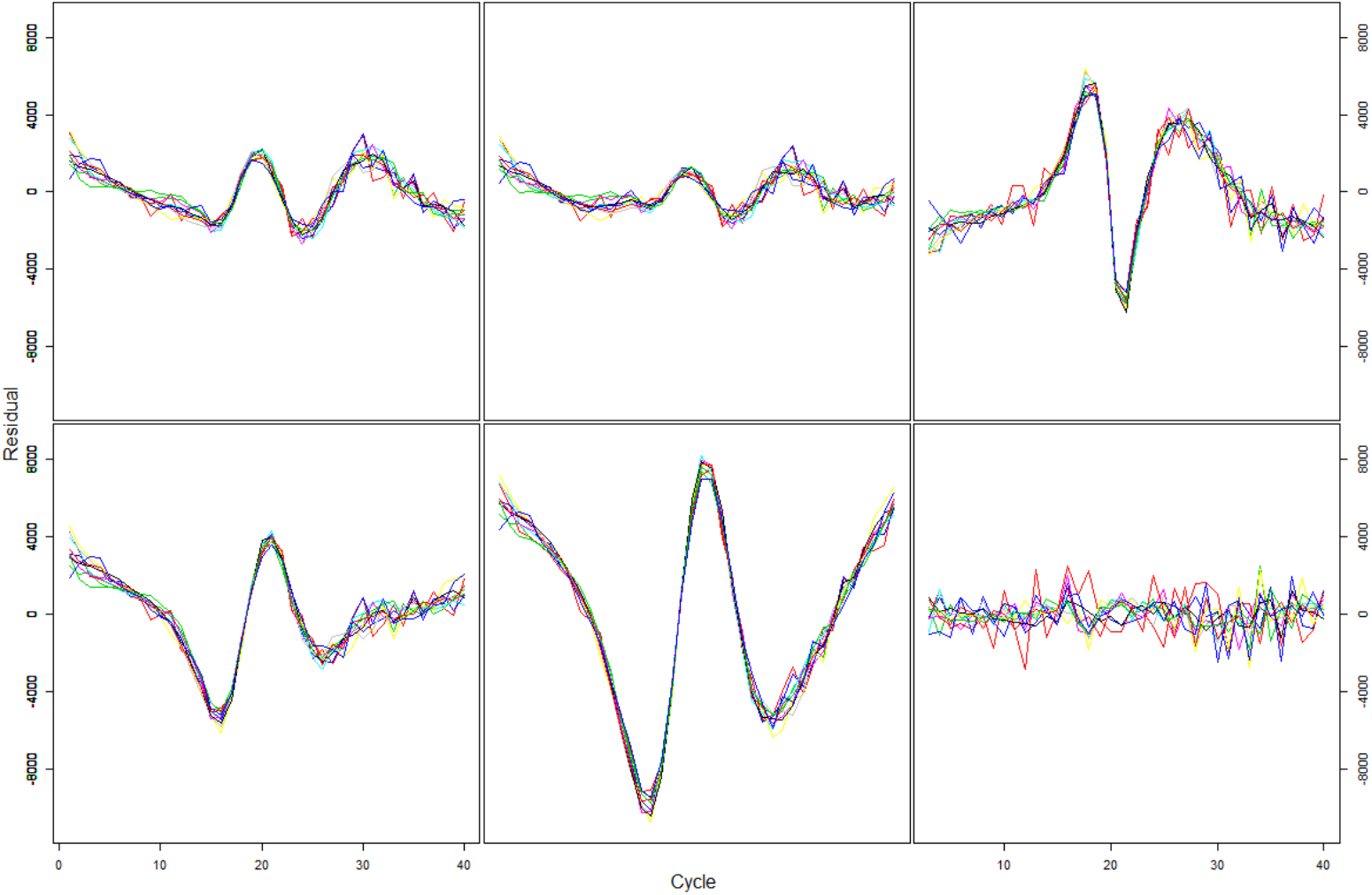
(Top left: 5-parameter log-logistic model (l5), Top middle: 5-parameter logistic model (b5), Bottom left: 4-parameter log-logistic model (l4), Bottom middle: 4-parameter logistic model (b4), Top right: LSTAR model with lag=1, Bottom right: LSTAR model with lag=2) All sigmoidal models and LSTAR model residuals are on the same y-axis scale. The residuals for each replicate curve vs. actual values for one set of replicates in GAPDH.SO are plotted by cycle progression. All subsets illustrate a between replicate non-random residuals problem for sigmoidal models. The LSTAR model with lag term of 1 also faces the same residuals problem, but LSTAR models with the lag term greater than 2, do not exhibit the problem and have random residuals.

When we apply the LSTAR model to see if it exhibits similar behavior, we also see that a lag term of 1, is unable to alleviate this problem (Figure 3 top right). An identical non-random residuals problem is present, similar to that of the sigmoidal models. However, when we expand our lag term to 2, the nonrandom residuals problem no longer exists, and the residuals appear to be random with mean zero and constant variance (Figure 3 bottom right). Additionally, the range for the residuals in the LSTAR model with a lag term of 2 is smaller than that of any of the other models.

### 3.2 RSS and Local RSS

To assess overall model fit, we compared the RSS and local RSS for the 5-parameter log-logistic model (l5) to the lag-2 LSTAR model using the full miRcomp data set. The l5 model was found to have higher RSS compared to the LSTAR model (Table 1). This suggests that from the perspective of the entire qPCR reaction, the LSTAR model is able to better capture the amplification curve than the l5 model. However, the mean local RSS for l5 is lower than for the LSTAR model, which suggests in the region around the Cq value, there may be a bias / variance trade-off between the l5 and LSTAR models. The local RSS for the LSTAR model on average, accounts for 855.02/6631 = 12.89% of the total RSS. In comparison, the local RSS for the l5 model accounts for 657.1/8603=7.64%. Of note, the local RSS calculation has significantly more NAs than the total RSS due to 1) NAs in the SDM method for Cq value calculation, and/or 2) Cq values estimates at the boundaries of the reaction for which a 4 cycle region is unavailable.

**Table 1.** Residual sum-of-squares and local residual sum-of-squares for the l5 and LSTAR model for all samples in miRcomp data set.

|  | Min | Median | Mean | Max | NAs |
| --- | --- | --- | --- | --- | --- |
| **RSS L5** | 512 | 4744 | 8603 | 30019173 | 989 |
| **Local RSS L5** | 0.1 | 294.5 | 657.1 | 1033850.6 | 2651 |
| **RSS LSTAR** | 322 | 5018 | 6631 | 5136253 | 1283 |
| **Local RSS LSTAR** | 0.03 | 563.92 | 855.02 | 100696.76 | 2669 |

### 3.3 Autocorrelation: Durbin-Watson and Ljung-Box Statistic

The Durbin-Watson test statistic allows for the detection of serial correlation amongst the residuals from the time-based progression in the amplification process. Under the null hypothesis, the residuals are serially uncorrelated, while the alternative hypothesis states that the residuals follow an autoregressive process with a time lag of one. When the 5-parameter sigmoidal model (l5) is applied to the miRcomp data set, we observe Durbin-Watson test statistics in the range of 0.05 to 3.00 (Figure 5), with the vast majority less than 2 indicating positive autocorrelation. The Ljung-Box test and lag 1 Pearson correlation show similar results (Supplementary Figure 5).

**Figure 5.**
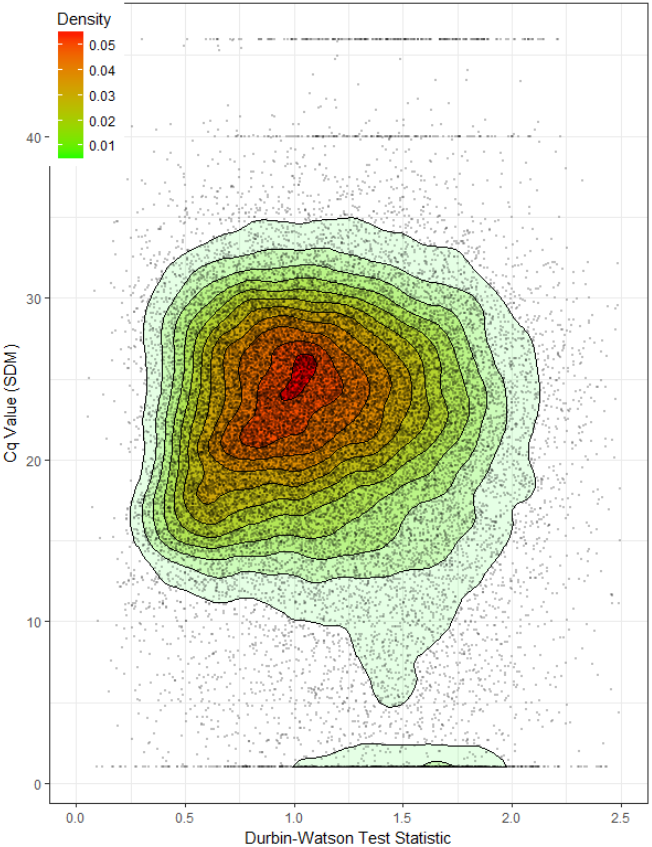
(Durbin-Watson Test Statistic for Residuals) Heat map with overlay of contour plot for the density distribution of Durbin-Watson test statistic and Cq value using the SDM. The 5-parameter log-logistic sigmoidal model was used. Here the values are from for all targets in miRcomp except the control targets, consisting of amplifications of 40 and 46 cycles. The range of the Durbin-Watson test statistic is from 0.05 to 3.00, with the majority less than 1. Cq values are consistently between cycles 15 to 30.

### 3.5 Categorization of qPCR Amplification

When estimating amplification curves from single qPCR reactions, we propose a collection of assessments based on both the observed data and the fitted values from a model. Specifically, we propose four general categories: good, poor, no signal, and failed reaction. We define a good fit as one in which the model has low RSS and minimal autocorrelation in the residuals. A poor fit may have a low RSS but substantial residual autocorrelation. In contrast the “no signal” and “failed reaction” categories are based primarily on the observed fluorescence data. “No signal” denotes a reaction for which the fluorescence values resemble random noise. A failed reaction occurs when the fluorescence peaks early and quickly decreases, suggesting an underlying issue with the amplification. Example of each of these categories are shown in Figure 5.

**Figure 5.**
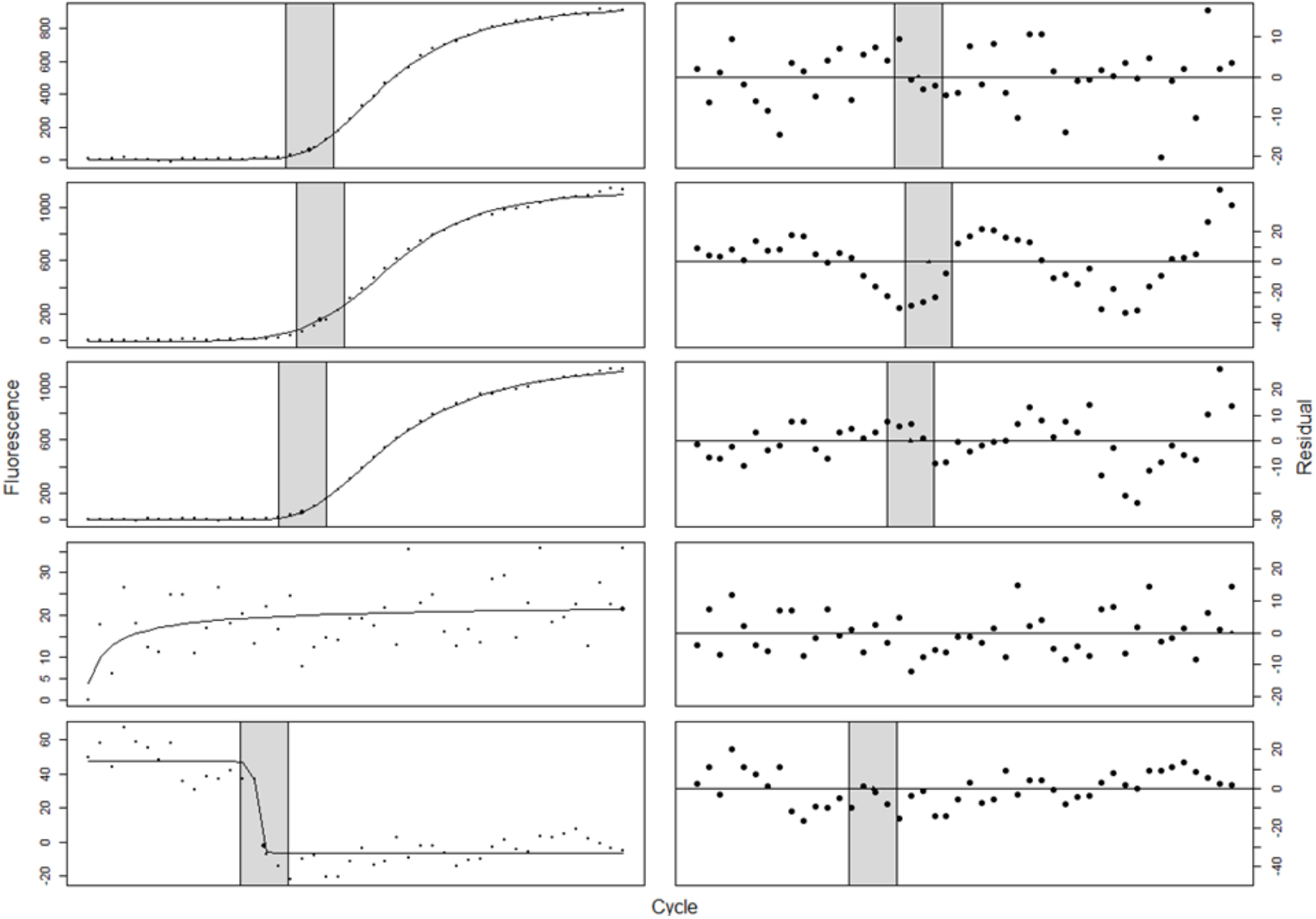
From top to bottom: (1-3) used target hsa-miR-23a_000399, while, (4-5) used target hsa-let-7c#_002405. All plots used a 5-parameter log-logistic model (l5), except (2) using a 4-parameter log-logistic model (l4). Here, (1) good signal fit for sample 8, replicate 4, (2) poor signal fit with symmetrical residual peaks for sample 9, replicate 4, (3) poor signal fit after asymmetrical factor included for sample 9, replicate 4, (4) non-signal fit for sample 3, replicate 4, and (5) failed-reaction signal fit for sample 9, replicate 2. Black circles (left column) or triangles (right column) represent the Cq values estimated by SDM. The shaded grey region denotes the region used to calculate the local RSS. There is no local RSS for the non-signal fit because the Cq value is nonsensical and appears at the boundary.

## Discussion

We have demonstrated that the sigmoidal curve fitting methods commonly used to analyze qPCR data deviate from the assumed model over the progression of the reaction. When estimating an average curve for several technical replicates, variation in the amount of the initial target molecule or in early amplifications appear to cause consistent differences in fluorescence across the PCR cycles. Furthermore, these differences are amplified during the exponential phase of the reaction. When fitting these sigmoidal models to individual qPCR reactions, we observed substantial autocorrelation in the residuals. This is perhaps not surprising given that these methods do not account for the dependence between consecutive cycles. As an alternative to sigmoidal models, we proposed a lag-2 Logistic Smooth Transition Autoregressive (LSTAR) model and showed that such a model addresses the limitations of the sigmoidal models by explicitly modeling the dependence between cycles and a gradual transition between amplification regimes.

While not explicitly addressed in this manuscript, the misspecification of the sigmoidal models likely effects the Cq estimates derived from the fitted models and thereby subsequent estimates of differential expression. While it is possible that Cq estimation via the second derivative maximum is relatively robust to the observed type of model misspecification, it should be noted that the introduction of an asymmetry parameter in the sigmoidal models (moving from 4-parameter to 5-parameter models) improved Cq estimation. This suggests similar improvements could be achieved by further improving the model fit via an LSTAR model or some alternative autoregressive modeling approach.

The presence of distinct phases in a PCR reaction has long been recognized; however, most current methods do not explicitly model these regimes. The standard view of a PCR reaction includes three phases: baseline, exponential, and plateau (Figure 1). However, the proposed LSTAR model captures the PCR dynamics using only two regimes: (1) a first phase in which the rate of the reaction is steadily increasing from an initial near negligible increase into the exponential phase up to the inflection point and (2) a second phase in which the rate of the reaction is steadily decreasing from the inflection point to the final plateau phase. By encoding a flexible transition between different stages of the amplification reaction, the LSTAR model better mimics the dynamics of the PCR reaction. Finally, the parameters of the LSTAR model are easily interpretable and can be linked back to specific aspects of the reaction.

While we have shown the potential of autoregressive models, especially the LSTAR model, to improve the analysis of qPCR data, additional performance assessments are warranted. Furthermore, we noted several technical challenges that may necessitate modifications of the LSTAR model. First, for curves with little noise, the grid search for the smoothing parameter often resulted in selection of the maximum value. One possible explanation for this could be these qPCR data already smoothly transition between the two regimes, making the smoothing parameter redundant. Second, our choice of estimating a Cq value from the LSTAR model by the change point between regimes is potentially suboptimal. Other parameters (or functions of several parameters) may provide better Cq estimates. Finally, while the LSTAR model achieved lower RSS than the 5-parameter sigmoidal model, the local RSS was higher for the LSTAR model. This is potential a result of differences in Cq estimation between the two methods rather than the models themselves, regardless further investigation is needed.

A conceptually appealing alternative to the sigmoidal and LSTAR models is a branching process model developed by Hanlon and Vidyashankar (2011). By targeting the stochastic process that governs the PCR reaction, they identify the variability between replicates as one of the primary causes of differences in efficiency estimates. To address this issue, they focused on isolating the exponential phase and treat the efficiency of the reaction as a random effect. This allows one to estimate the between reaction variability in efficiency. Incorporating random effects into a branching process model yields a probabilistic data generating model that reflects the probabilistic nature of PCR amplifications. In contrast to sigmoidal and LSTAR models, which model the entire PCR reaction, the branching process model uses only the exponential phase of the reaction. However, currently this method lacks publicly available software and requires identification of the exponential phase, for which we are unaware of any available methods. While these issues prevented us from evaluating this branching process model in this manuscript, we are intrigued by the potential of such methods to model qPCR data.

## Supplementary Materials

### Supplemental Note

The general formula for the STAR model with the additional parameter for embedding dimension, m, is:

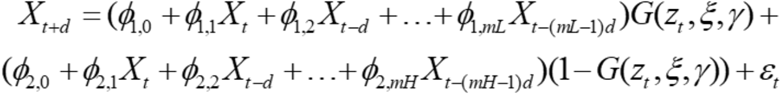

Since qPCR does not contain cyclical trends, we omit the dimension parameter. We consider a model in which the delay parameter and embedding dimension both equal to one. This yields the following equations:

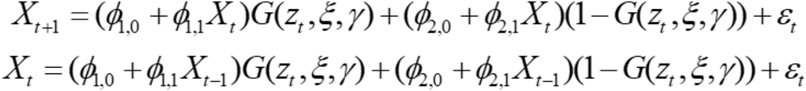

Next, we consider a model in which the delay parameter equals one and the embedding dimension equals two. This yields the following model equations:

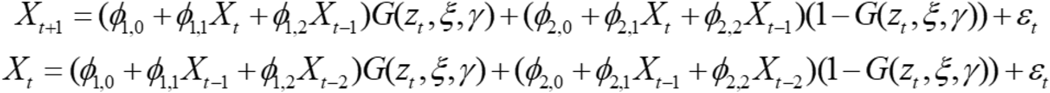

Note that by altering the embedding dimension, we are able to obtain STAR models with lag terms that are multipliers of one.

**Supplementary Figure 1:**
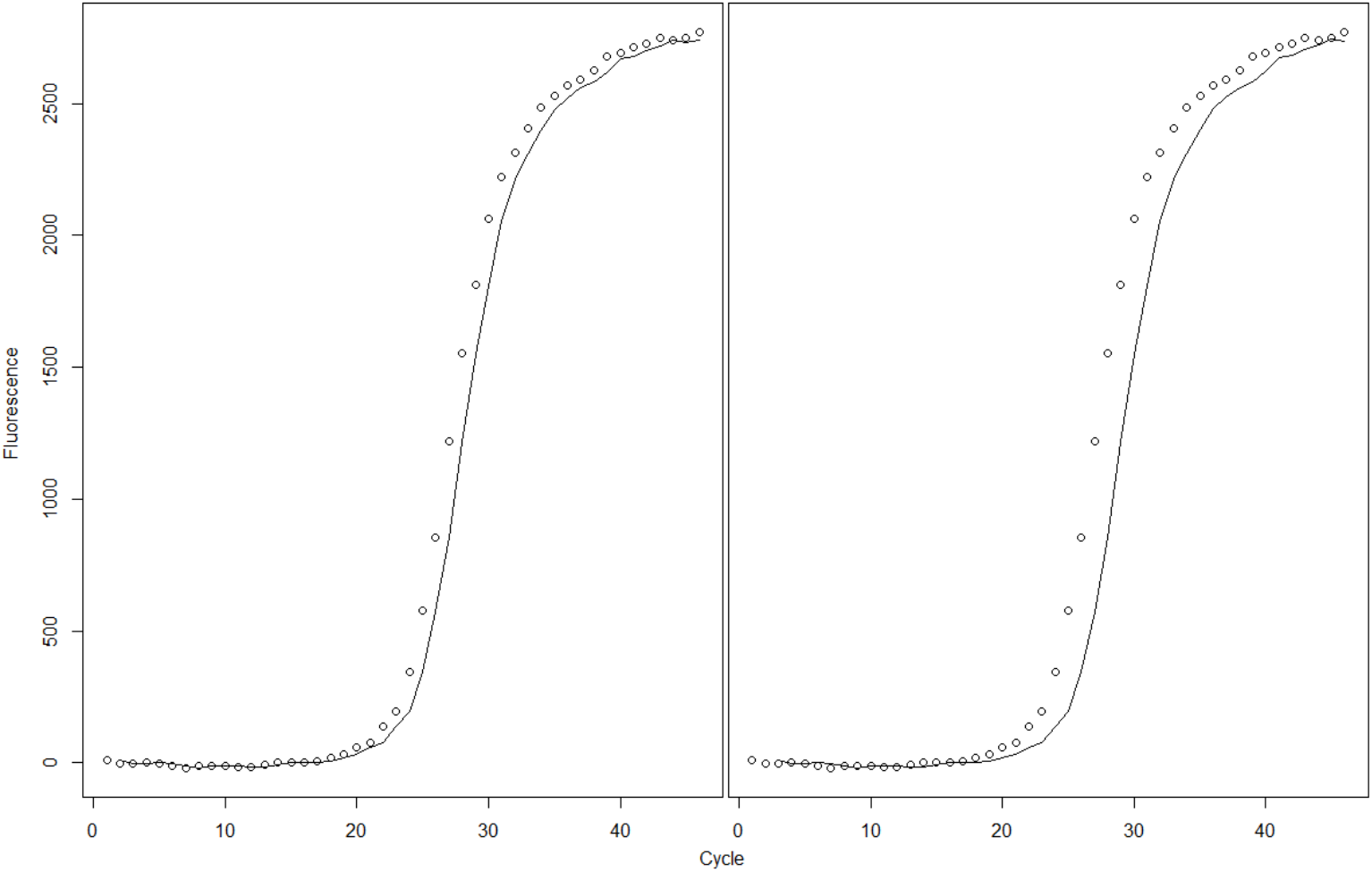
Autoregressive models on miRcomp target hsa-miR-520e_001119, replicate E1. (Left: AR(1) with coefficient of 0.996, (2) AR(2) with coefficients of 1.978, and −0.981. Both illustrate the problems with using a direct autoregressive model that is linearly dependent on its own lagged series. The values will of the current cycle will not be able to adjust accordingly to any shifts in phase, because there are no regime-switching properties included.

**Supplementary Figure 2:**
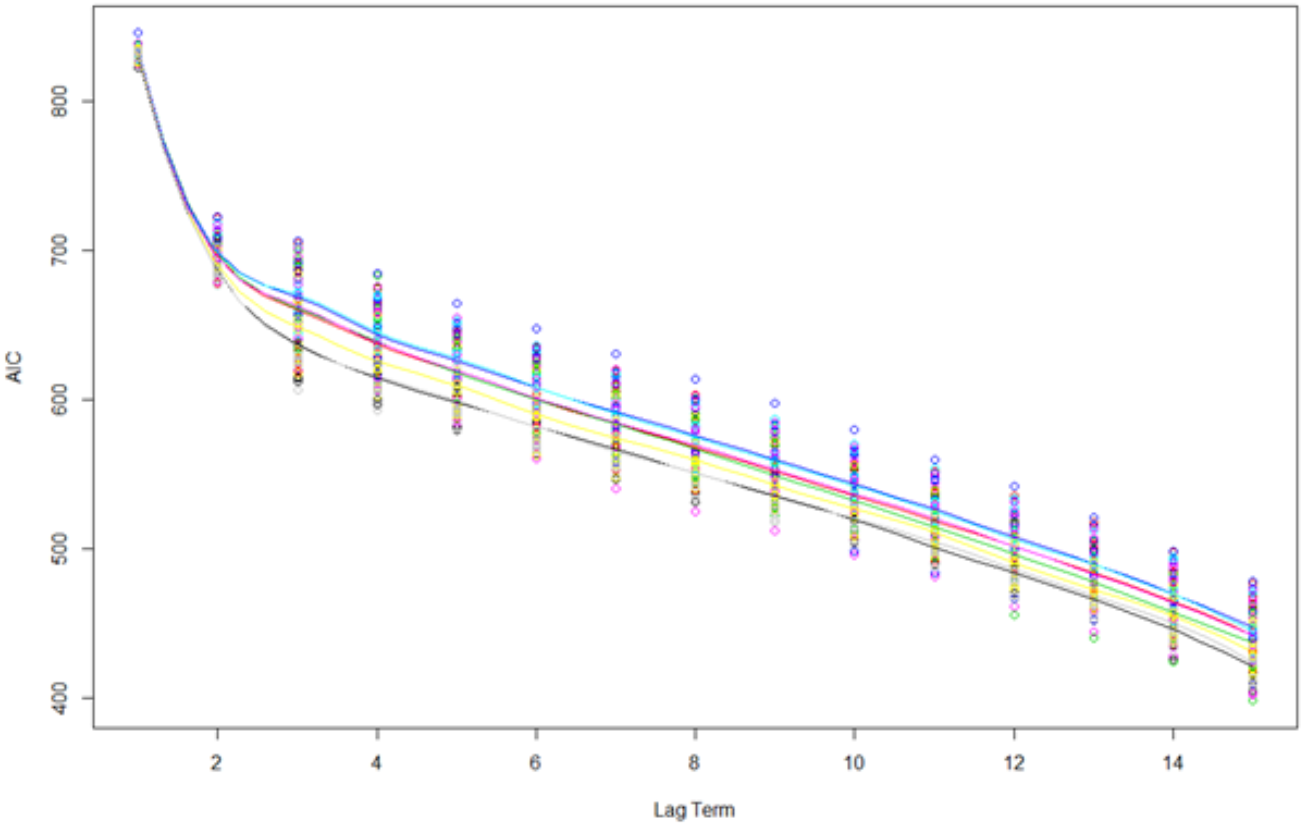
AIC for a dynamic linear model with different lag terms in GAPDH data set, fitted with smoothing curve. There are improvements in AIC with the inclusion of additional lag term, however, none as significant as the inclusion of lag term 2. Colored points denote all replicates of subsets in GAPDH. The AIC values here provide compelling evidence for the use of a lag of 2 in the STAR model.

**Supplementary Figure 3:**
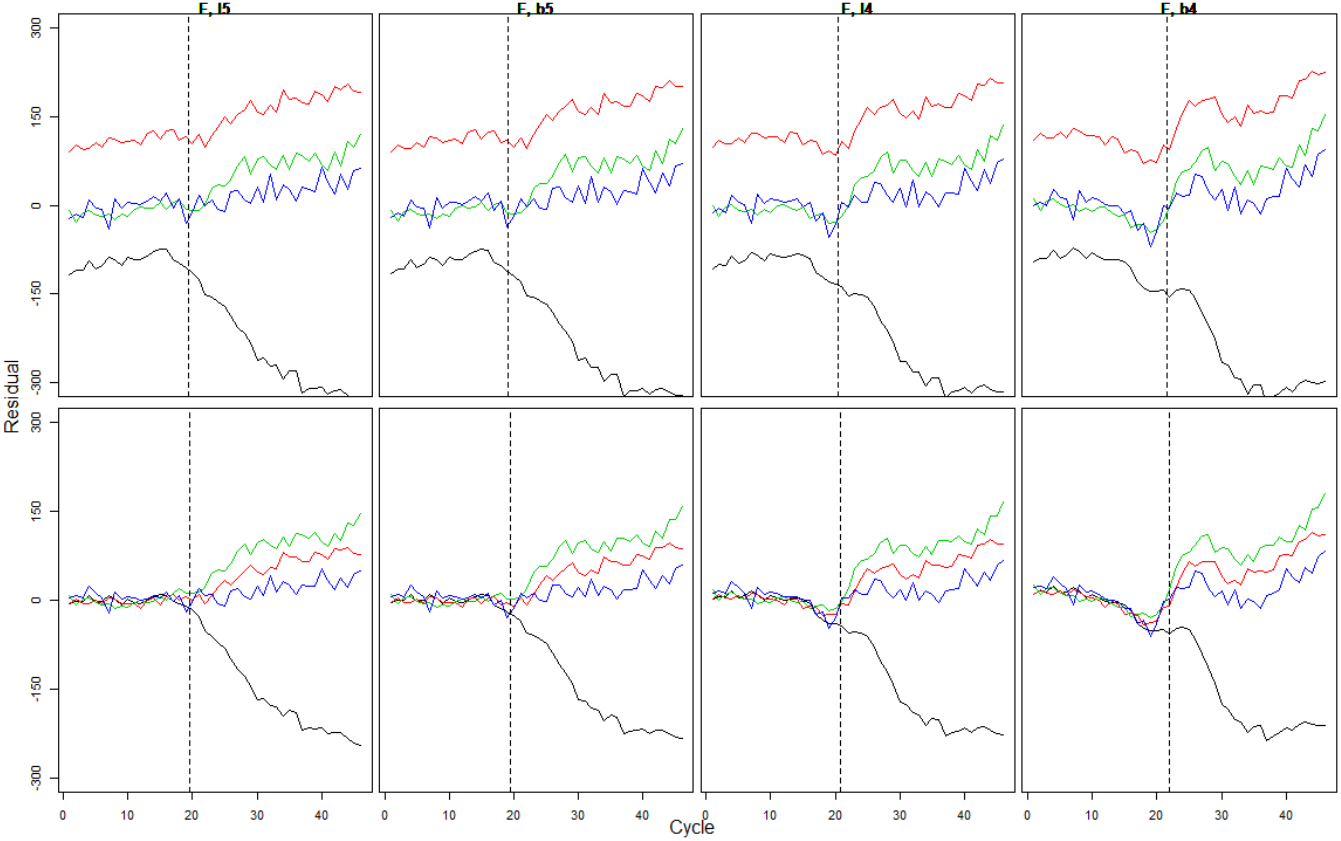
(Left to right: 5-parameter log-logistic model (l5), 5-parameter logistic model (b5), 4-parameter log-logistic model (l4), 4-parameter logistic model (b4). Top row: raw fluorescence, bottom row: baseline subtracted fluorescence) Between replicate residuals for subset F of miRcomp target hsa-miR-23a_000399. This data consists considerably more noise in its amplification process, but panels provide similar patterns to that of GAPDH for both non-baseline subtracted and baseline subtracted fluorescence. Specifically, an amplified affect as the cycle progress after the dashed line (Cq value). The baseline has residuals around 0, after subtracting the fluorescence values by the minimum values of each replicates, but the problem is not fixed.

**Supplementary Figure 4:**
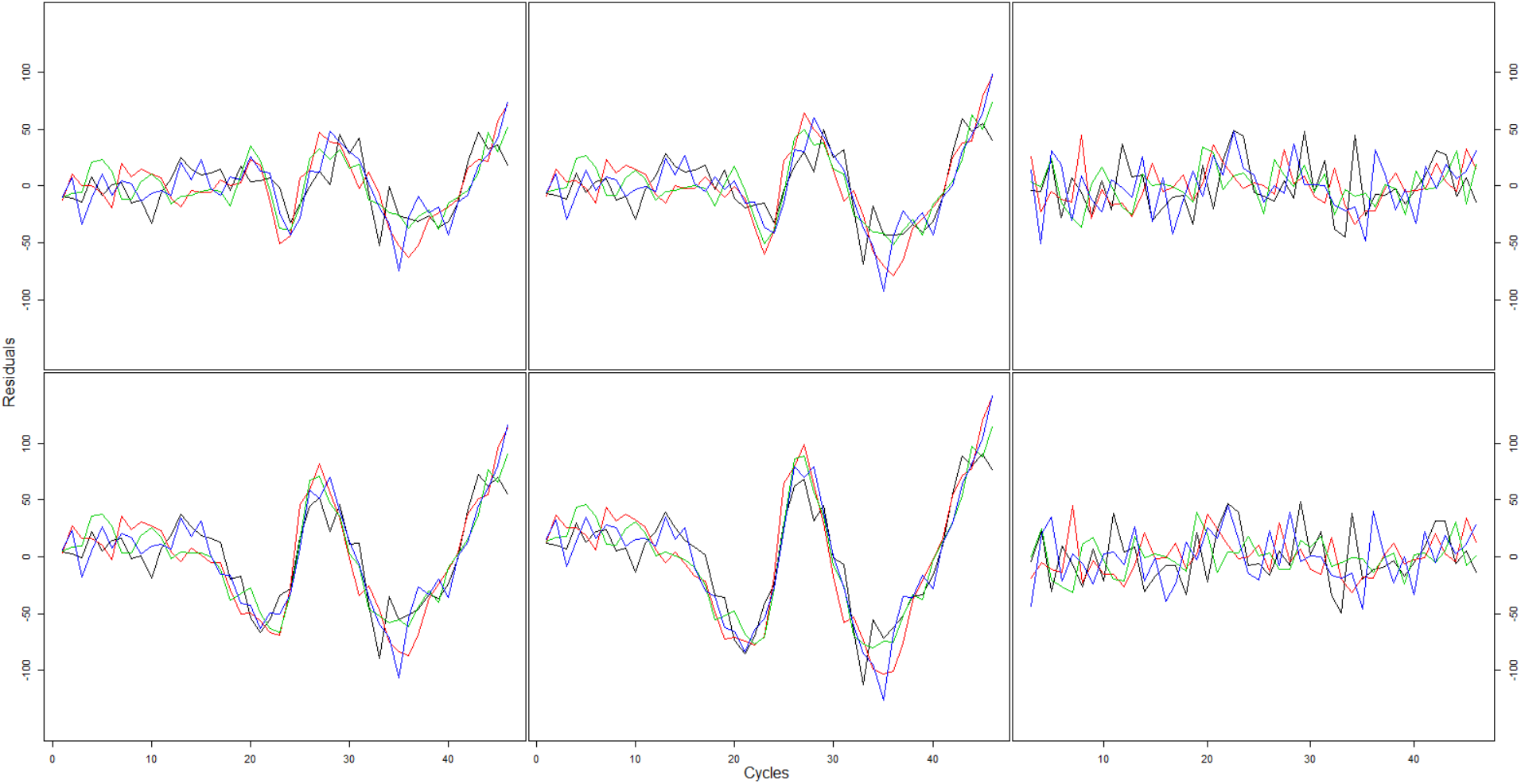
(Top left: 5-parameter log-logistic model (l5), Top middle: 5-parameter logistic model (b5), Bottom left: 4-parameter log-logistic model (l4), Bottom middle: 4-parameter logistic model (b4), Top right: LSTAR model with lag=1, Bottom right: LSTAR model with lag=2) The residuals for each replicate curve vs. actual baseline-subtracted values within subset H for miRcomp target hsa-miR-500_002428 are plotted by cycle progression. All subsets illustrate a between replicate non-random residuals problem for sigmoidal models. The non-random residual problems does not seem to be present in the LSTAR models for both lag 1 and 2.

**Supplementary Figure 5:**
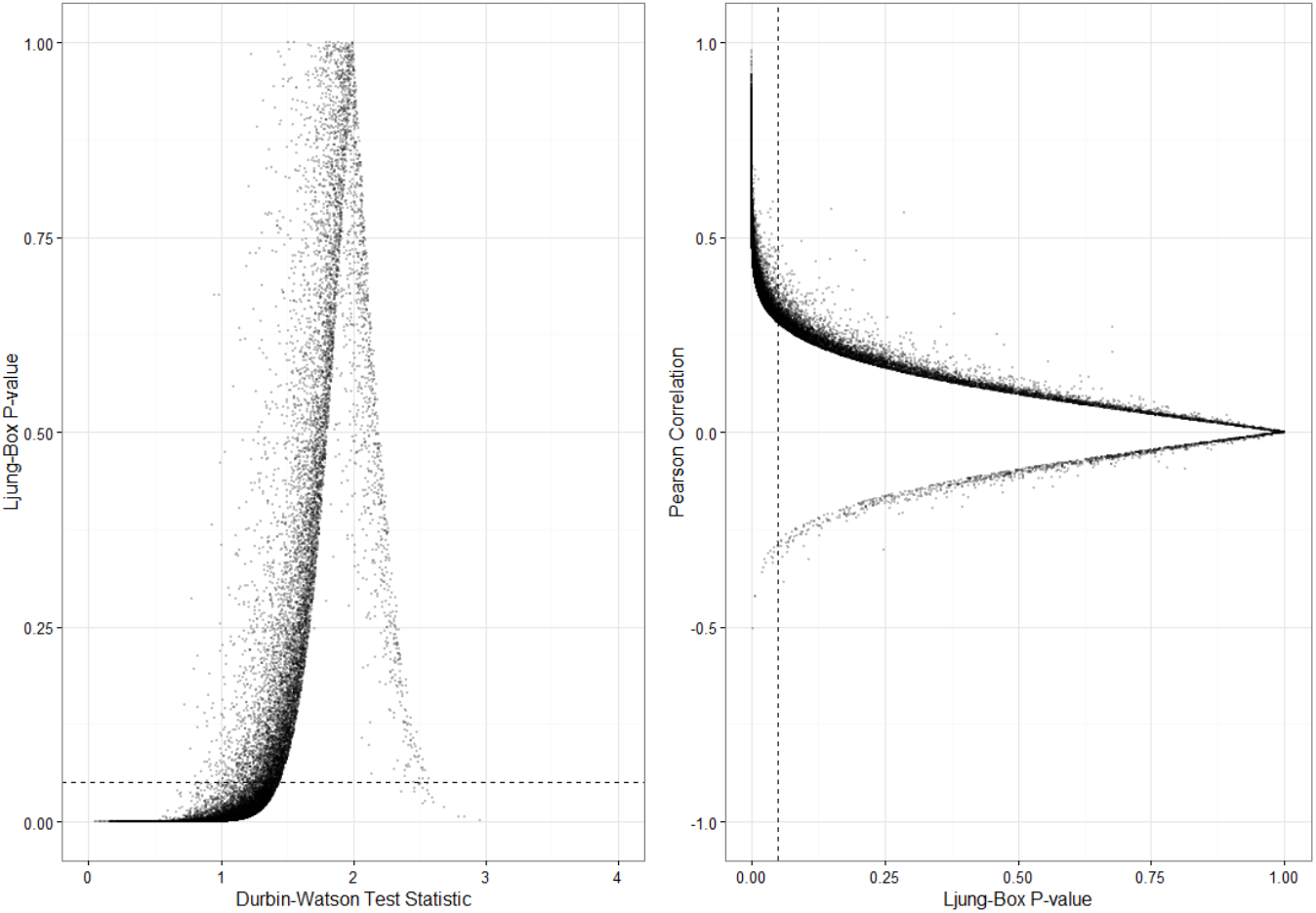
Though the Durbin-Watson, Ljung-Box, and Pearson correlation all provide estimates to test if the residuals exhibit autocorrelation, each has different estimates. The test-statistic for the Durbin-Watson, Ljung-Box p-value, and Pearson correlation are related in different ways. We show that after fitting the miRcomp data with a 5-parameter log-logistic sigmoidal model, the results of a statistically significant Ljung-Box p-value corresponds to a Durbin-Watson test-statistic of approximately 1.4 and below, and a Pearson correlation of at least 0.3 (absolute value).

## Bibliography

Bar, Tzachi, Mikael Kubista, and Ales Tichopad. “Validation of kinetics similarity in qPCR.” Nucleic acids research 40.4 (2011): 1395–1406.

Bustin, Stephen A., et al. “The MIQE guidelines: minimum information for publication of quantitative real-time PCR experiments.” Clinical chemistry 55.4 (2009): 611–622.

Chen, Yanguang. “Spatial autocorrelation approaches to testing residuals from least squares regression.” PloS one 11.1 (2016): e0146865.

Hanlon, Bret, and Anand N. Vidyashankar. “Inference for quantitation parameters in polymerase chain reactions via branching processes with random effects.” Journal of the American Statistical Association 106.494 (2011): 525–533.

Lievens, Antoon, et al. “Enhanced analysis of real-time PCR data by using a variable efficiency model: FPK-PCR.” Nucleic acids research 40.2 (2011): e10–e10.

Lim, K. S., and H. Tong. “Threshold autoregressions, limit cycles, and data.” Journal of the Royal Statistical Society B, 42 (1980): 245–92.

Livak, Kenneth J., and Thomas D. Schmittgen. “Analysis of relative gene expression data using real-time quantitative PCR and the 2-ΔΔCT method.” Methods 25.4 (2001): 402–408.

Ljung, Greta M., and George EP Box. “On a measure of lack of fit in time series models.” Biometrika 65.2 (1978): 297–303.

McCall, Matthew N., et al. “A benchmark for microRNA quantification algorithms using the OpenArray platform.” BMC bioinformatics 17.1 (2016): 138.

Rutledge, Robert G. “A Java program for LRE-based real-time qPCR that enables large-scale absolute quantification.” PLoS One 6.3 (2011): e17636.

Spiess, Andrej-Nikolai, Caroline Feig, and Christian Ritz. “Highly accurate sigmoidal fitting of real-time PCR data by introducing a parameter for asymmetry.” BMC bioinformatics 9.1 (2008): 221.

Teräsvirta, Timo. “Specification, estimation, and evaluation of smooth transition autoregressive models.” Journal of the American Statistical Association 89.425 (1994): 208–218.

Tong, Howell. “On a threshold model.” In: Chen, C, (ed.) Pattern Recognition and Signal Processing. NATO ASI Series E: Applied Sc. (29). Sijthoff & Noordhoff, Netherlands (1978): 575–586.

Tong, Howell. “Non-linear time series. A dynamical system approach.” Oxford Statistical Science Series, Oxford: Clarendon Press, 1990 (1990).

